# Physicochemical Properties of Chromosomes in Live Cells Characterized by Label-Free Imaging and Fluorescence Correlation Spectroscopy

**DOI:** 10.1101/533596

**Authors:** Tae-Keun Kim, Byong-Wook Lee, Fumihiko Fujii, Kee-Hang Lee, YongKeun Park, Jun Ki Kim, Chan-Gi Pack

**Author notes:** **Correspondence:** Jun Ki Kim, Chan-Gi Pack. T.-K. K. and B.-W. L. contributed equally to this work.

## Abstract

The cell nucleus is a three-dimensional, dynamic organelle that is organized into many subnuclear bodies, such as chromatin and nucleoli. The structure and function of these bodies is maintained by diffusion and interactions between related factors as well as dynamic and structural changes. Recent studies using fluorescent microscopic techniques suggest that protein factors can access and are freely mobile in mitotic chromosomes, despite their densely packed structure. However, the physicochemical properties of the chromosome itself during cell division are not yet fully understood. Physical parameters, such as the refractive index (RI), volume of the mitotic chromosome, and diffusion coefficients of fluorescent probes inside the chromosome were quantified using an approach combining label-free optical diffraction tomography with complementary confocal laser scanning microscopy and fluorescence correlation spectroscopy. Variance in these parameters correlated among various osmotic conditions, suggesting that changes in RI are consistent with those in the diffusion coefficient for mitotic chromosomes and cytosol. Serial RI tomography images of chromosomes in live cells during mitosis were compared with three-dimensional confocal micrographs to demonstrate that compaction and decompaction of chromosomes induced by osmotic change were characterized by linked changes in chromosome RI, volume, and the mobility of fluorescent proteins.

## 1 Introduction

The cell nucleus is dynamically and three-dimensionally organized into subnuclear bodies including chromatin, chromosomes, and nucleoli, and contains a high density of macromolecules. In particular,the long string of genomic DNA is organized in nuclei and mitotic chromosomes to make genomic information accessible for the many physiological functions of cells. While it remains unclear how protein factors find and interact with their targets, the structure and function of cellular bodies are likely maintained by mobility and interaction with related factors during the cell cycle. Previous studies using biophysical methods such as fluorescence correlation spectroscopy (FCS) demonstrated that various probe molecules freely diffuse in the nucleus and various subnuclear bodies, such as chromatin, chromosomes, and nucleoli (Hihara et al., 2012; Pack et al., 2006; Park et al., 2015; Wachsmuth et al., 2008). FCS based on confocal laser scanning microscopy (CLSM) was applied as a useful and highly sensitive technique for quantitatively assessing molecular concentration and diffusion of fluorescent probe in aqueous solutions and living cells (Eigen and Rigler, 1994; Erdel et al., 2011; Magde et al., 1974; Pack et al., 2006).

The cellular microenvironment and molecular crowding are key physicochemical characteristics of cellular bodies with a high macromolecular content (Gnutt et al., 2015; Hancock, 2004; Matera et al., 2009; Richter et al., 2007; Richter et al., 2008), and diffusion coefficients (*D*s) of fluorescent probe molecules such as green fluorescent protein (GFP) in live cells have been systematically characterized to investigate these traits (Finan et al., 2011; Hihara et al., 2012; Konopka et al., 2009; Mika et al., 2014; Park et al., 2015; Schavemaker et al., 2018; Verkman, 2002). In particular, observations using FCS suggested that heterochromatin and mitotic chromosomes are accessible to protein factors with a molecular size up to 150 kDa, despite their densely packed structure (Hihara et al., 2012). It was also suggested that nucleosomes as well as probe proteins are freely mobile in these bodies. However, the physicochemical properties of mitotic chromosomes under various physiological conditions are not yet fully understood. Since FCS is highly sensitive and only requires a small detection volume, it is well-suited to measuring the diffusional mobility of probe molecules in very small regions that comprise subnuclear compartments in living cells. However, FCS measurements of large areas are time-consuming and too inefficient to allow simultaneous volumetric measurement of mobile bodies such as the mitotic chromosome. Moreover, the phototoxic and bleaching effects of fluorescence methods such as confocal microscopy and FCS must be also carefully considered when attempting to obtain reliable information from live cells, especially mitotic cells.

In the present study, we used three complementary methods in a single approach, combining the label-free quantitative phase imaging (QPI) method with CLSM and confocal-based fluorescence correlation spectroscopy to compensate for the limitations of the fluorescence method, such as phototoxicity, long scanning time for three-dimensional (3D) imaging, and time-consuming measurement of FCS. Recently, several label-free QPI methods, such as optical diffraction tomography (ODT), were identified as promising methods for high-speed live cell imaging (Brunsting and Mullaney, 1974; Choi et al., 2007; Kemper et al., 2011; Kim et al., 2013; Kim et al., 2014). Furthermore, as low light intensities are required for object illumination, ODT minimizes the photostress on the biological sample, making it suitable for noninvasive measurement of live cells during mitosis. Furthermore, ODT provides information on absolute biophysical parameters such as volume and refractive index (RI) (Liu et al., 2016). Since the RI is generally proportional to the concentration of organic solutes, which, in turn, is related to the viscosity of aqueous solutions (Sung et al., 2012), it is expected that FCS and ODT will be complementary to each other.

ODT is an interferometric microcopy technique that acquires 3D RI tomograms of cells and tissues without prior preparation or labeling. Therefore, ODT microscopy can observe unfixed cells and unlabeled, living cells without fluorescent protein expression or immunofluorescence. Moreover, it is fast and can acquire one 3D RI tomogram in <1s (Park et al., 2010). ODT has been commercialized and is easy to use for label-free imaging of live cells.

Indian Muntjac (DM) cells are an ideal cell line to visualize diffusion of fluorescent proteins through the cell chromosome. Among mammals, the male DM cells have 2n = 7 diploid chromosomes that are large compared with common culture cells such as HeLa cells. DM cells expressing H2B-monomer RFP (H2B-mRFP) and monomer GFP (mGFP) were subjected to fluorescence imaging methods (Hihara et al., 2012) to enable a direct comparison between ODT and fluorescent confocal 3D micrographs of the mitotic chromosomes. In addition, mGFP may be used as a fluorescent probe to quantify *D* and local viscosity in the mitotic chromosome.

In the present study, we demonstrated the application of our method for quantification of several physical parameters of the mitotic chromosome, including volume, RI, and diffusion coefficient (*D*) for fluorescent probes inside the chromosome. We successfully acquired 3D RI images of mitotic chromosomes of DM cells and combined analyses with CLSM and FCS. Due to the small number and large size of the chromosomes, tomographic RI images of DM cells showed a clear structure of chromatids compared with that of HeLa cells. The RI tomographic images of chromosome of DM cells were consistent with the fluorescence images of these cells expressing H2B-mRFP. Moreover, we quantitatively compared the RI values of chromosomes, molecular accessibility, and *D* value of mGFP probe inside mitotic chromosome under various osmotic conditions. Confocal and RI images showed that chromosomes in the mitotic cells were significantly compacted by treatment with a solution of high osmolality, and indicated that hypotonic stress induced decompaction of the chromosomes. This compaction and decompaction of chromosomes characterized by change in RI was consistent with the changes observed in *D* value of the mGFP probes.

## 2 Materials and Methods

### 2.1 Cells

DM cells stably expressing H2B-mRFP and mGFP (Hihara et al., 2012; Pack et al., 2006) were kindly provided by Dr. Maeshima (National Institute of Genetics, Japan). DM cells were cultured at 37°C in 5% CO_2_ in Dulbecco’s modified Eagle’s medium (DMEM) supplemented with 20% fetal bovine serum (FBS), 100 U/mL penicillin, and 100 U/mL streptomycin. HeLa cells were cultured at 37°C in 5% CO2 in DMEM supplemented with 10% FBS, 100 U/mL penicillin, and 100 U/mL streptomycin. For live cell microscopic analysis, stable DM cells were plated in tomo-dishes (Tomcube, Korea) or Lab-Tek 8-well chambered coverglass (Nunc, USA). For observation of HeLa cells expressing H2B-mRFP, cells were transiently transfected using lipofectamine 3000 (Thermo Fisher Scientific, USA).

To investigate the influence of osmotic stress, DM cells were incubated with either phosphate-buffered saline (PBS) for hypotonic conditions, or phenol-free DMEM supplemented with 20% FBS and 0.2 M sucrose for hypertonic condition. Media were prewarmed to 37°C prior to application. For HeLa cells, PBS or phenol-free DMEM supplemented with 10% FBS and 0.2 M sucrose was applied prior to imaging. Dulbecco’s PBS (Biowest, France) was used for DM and HeLa cells.

### RI of Solutions

The RI of glycerol–water solutions and culture media was measured using a refractometer with temperature control from 15°C to 60°C (Abbemat 3200; Anton Paar GmbH, Austria) (Table 1). Glycerol solutions with purity >99.5% (Biosesang, Korea) were used at concentrations of 10%, 20%, 30%, 40%, 50%, 60%, and 70% (w/v) to determine the relationship between viscosity (Weast, 1988) and RI.

**Table 1.** Measured refractive indices of culture media and solutions at 25°C and 37°C.

|  | **DMEM<sup>a</sup>**<br>**(20%**<br>**FBS)** | **0.2 M<sup>b</sup>**<br>**sucrose**<br>**(20%**<br>**FBS)** | **DMEM**<br>**(10% FBS)** | **0.2M**<br>**sucrose**<br>**(10%**<br>**FBS)** | **PBS only** | **Distilled**<br>**water**<br>**(reference)** |
| --- | --- | --- | --- | --- | --- | --- |
| RI (25°C) | 1.3365 | 1.3456 | 1.3358 | 1.3447 | 1.3349 | 1.3325 |
| RI (37°C) | 1.3350 | 1.3441 | 1.3342 | 1.3431 | 1.3336 | 1.3310 |

<sup>a</sup> DMEM containing 20% FBS and <sup>b</sup> DMEM containing 20% FBS and 0.2M sucrose were used as media for Indian Muntjac (DM) cells. DMEM containing 10% FBS was used for HeLa cells. Dulbecco's PBS was used for both DM and HeLa cells. The refractive index (RI) of distilled water is shown as a reference. RI was measured at 488 nm.

### 2.2 ODT

ODT was performed as described previously (Park et al., 2010). A Mach–Zehnder interferometric microscope was used to reconstruct the cell 3D RI tomograms (Mourao et al., 2014). This commercial microscope (HT-1H, Tomocube Inc., Korea) consists of an illumination/sample modulation unit and an optical field recording unit. Optical interference is used to capture amplitude and phase information from light transiting the sample (Kim et al., 2016).

### 2.3 Image rendering and volume calculation

The RI isosurfaces were rendered using commercial software (TomoStudio, Tomocube Inc., Korea). Volume was calculated using commercial software from Tomocube Inc. (Chromosome analysis, Tomocube Inc., Korea).

### 2.4 CLSM

Fluorescence microscopy using live cells was performed using an inverted confocal laser scanning microscope (LSM780; Carl Zeiss, Germany). For 3D and time-lapse images, fluorescently stained DM or HeLa cells were excited at a wavelength of 561 nm and the emission signal band was detected at 570–630 nm. The interval for time-lapse imaging was 3 min and z-stack images were taken at 0.5-µm intervals using a 5× magnification objective lens (C-Apochromat, 63 × /1.2NA). All live cell measurements were performed at 37°C in 5% CO2 culture conditions. Fluorescence images were processed using software (Zen 2012 SP5; Carl Zeiss, Germany) installed on a LSM780 confocal microscope system for 3D reconstruction or intensity profile analyses. Images taken using the confocal microscope were also 3D rendered using IMARIS 8.1.2 software (Bitplane, USA) and used for volume analysis. Where necessary, the surface function of the IMARIS software was used.

For fixed cell observation, cells were first cultured in a confocal microscope dish and fixed for 5 min with 1% paraformaldehyde and 1% glutaraldehyde in 0.1 M PBS. Fixed cells were stained for 5 min with DAPI. The photography conditions were enlarged five-fold using a 63× C-Apochromat lens, z-position thickness in 0.50 µm size unit, and 3D rendered using Zen 2012 SP5 or IMARIS software. The 3D panoramic images were processed using IMARIS. All CLSM observations were performed at 25°C.

### 2.5 FCS

Confocal imaging for FCS measurement was performed using an LSM780 confocal microscope system. mGFP was excited at 488 nm using a CW Ar^+^ laser through a water-immersion objective (C-Apochromat, 40×, 1.2 NA; Carl Zeiss). H2B-monomer RFP (mRFP) was imaged using a 561 nm DPSS laser. To avoid bleed-through effects in double-scanning experiments, mGFP and mRFP were scanned independently in multitracking mode, and mRFP was only scanned for FCS measurement. All CLSM observations and corresponding FCS measurements were performed at 25°C as described previously (Hihara et al., 2012; Kang et al., 2018).

FCS measurements were all performed at 25°C using an LSM780 confocal microscope (Carl Zeiss, Germary) as described previously (Hihara et al., 2012; Kang et al., 2018). mGFP was excited with minimized power at 488 nm with a continuous wave Ar^+^ laser through a water-immersion objective (C-Apochromat 40×/1.2NA; Carl Zeiss). H2B-mRFP was imaged using a continuous wave 561 nm laser. Image detection was performed using a GaAsP detector (Quasar; Carl Zeiss). All fluorescence autocorrelation functions (FAFs) were measured for 5 s at least three times with 2-s intervals to avoid nonstationary fluorescent fluctuations due to drift of the targeted chromosome over the measurement duration. To avoid bleed-through effects in double-scanning experiments, mGFP and mRFP were scanned independently using a multitracking mode, and only mRFP was scanned for FCS measurement.

Data analysis for FCS measurement was performed as previously described (Hihara et al., 2012; Pack et al., 2006). To obtain the diffusion time, FAFs [G (τ)] of the measurements were fitted using the following one-component model with or without a triplet term:

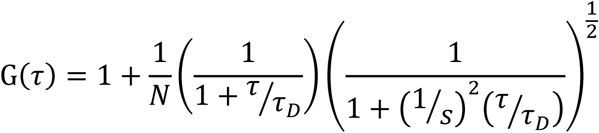

where *N* is the number of molecules in the detection volume, τ*_D_* is the correlation time, *w* and *z* are width and axial length of the detection volume, respectively, and *s* is the structure parameter *z*/*w*. Diffusion times had the following relationship to the diffusion coefficient:

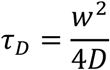

The diffusion coefficient of mGFP (*D*_mGFP_) was calculated from the reported value of the diffusion coefficient of rhodamine 6G (*D*_Rh6G_ = 280 μm^2^/s) and the corresponding measured diffusion times of Rh6G (*τ*_Rh6G_) and mGFP (*τ*_mGFP_) were as follows (Pack et al., 2006):

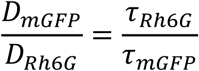

All FAF curves for DM cell FCS measurements fitted well to a one-component model, using Zen 2012 SP5 software, as demonstrated previously (Hihara et al., 2012). Glycerol–water solutions containing Rh6G were measured and the FAFs were analyzed to evaluate the *D* values and the corresponding viscosity (Pack et al., 2006).

### 2.6 Statistical methods

Student’s *t* tests were performed for comparison of mean values to determine significance (Origin v.8.5, Northampton, MA, USA). All *p*-values <0.05 were considered significant.

## 3 Results and Discussion

### 3.1 Comparisons between ODT and CLSM 3D images of DM cell chromosomes

To demonstrate how RI images of mitotic chromosomes were accurately constructed by ODT measurement, live DM cells expressing H2B-mRFP and mGFP were measured by ODT. Due to the small number and large size of the DM chromosomes, it was expected that tomographic RI images of DM cells would have a clear chromosomal structure that would enable the characteristic properties of chromosomes to be determined during mitosis. For each measurement of ODT and CLSM, label-free and 3D RI images of the mitotic chromosomes of a DM cell obtained from time-lapse ODT measurements and analysis were compared with fluorescent 3D images of H2B-mRFP during mitosis of other DM cell obtained by confocal microscopy (Fig. 1, Movie S1, S2). Figure 1A shows four tomographic RI images of a DM cell chromosome from prophase to telophase during mitosis, whereas Fig. 1B shows corresponding representative fluorescence images from H2B-mRFP expressed DM cells. Using time-lapse and 3D confocal imaging (Fig. 1B, Fig. S1a, Movie S3), the number of diploid chromosomes in the DM cells were counted and found to be seven, which is consistent with the known number of chromatids in male DM cells. In contrast, it was difficult to clearly image and count the number of chromatids in live HeLa cells using time-lapse and 3D observations of CLSM and ODT, respectively (Fig. S1b, Movie S4, S5). RI images of the DM cell chromosomes in prophase were not clear compared with those of other mitotic phases, and compared with the fluorescence images obtained using CLSM. The number of diploid chromosomes detected by CLSM could not be counted precisely from the RI images of the cells in metaphase or anaphase.

**Figure 1.**
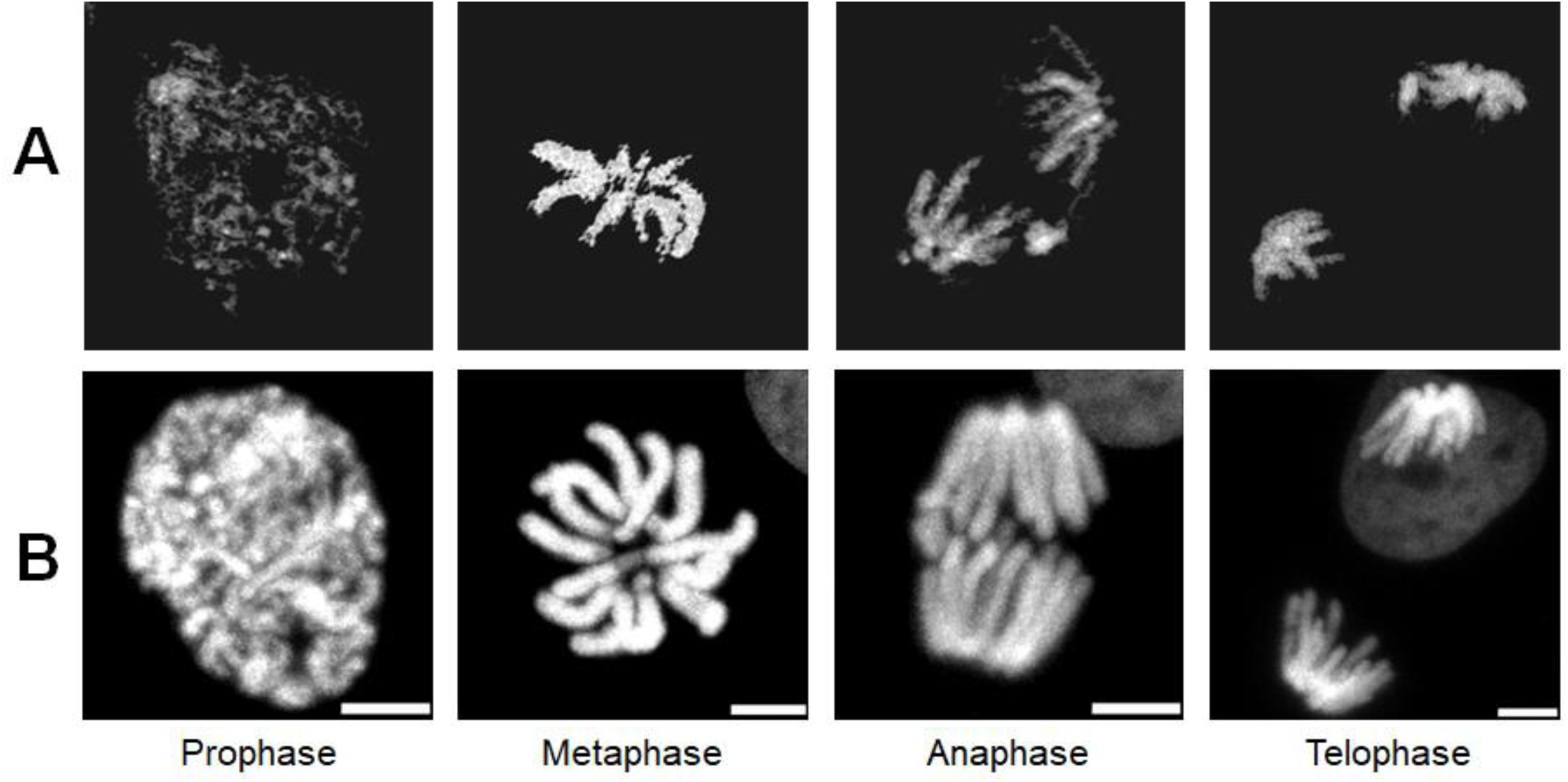
Comparison of 3D confocal fluorescence image of H2B-mRFP with 3D RI image of chromosome in mitosis. (**A**) Four RI images of a DM cell stably expressing H2B-mRFP and mGFP from a series of time-lapse observation during mitosis. For clarity, only high and continuous RI regions of chromosome in mitotic cytosol are shown (see also Supplementary Movie 1). (**B**) Four fluorescence images of H2B-mRFP from a series of time-lapse and 3D observations depicting mitosis in a DM cell stably expressing H2B-mRFP and mGFP. For clarity, images from mGFP channels are not shown. Scale bar represents 5 μm.

Nevertheless, the resulting images showed that ODT measurement allowed successful reconstruction of dynamic structural changes in the mitotic chromosomes of DM cells from metaphase to telophase.

### 3.2 Quantification of the RI of mitotic chromosomes at different osmotic conditions

To investigate the influence of osmotic changes on the structure and physicochemical properties of mitotic chromosomes in live DM cells, the physiological state of cells was changed using culture media of varying osmotic pressure (Table 1, see also Materials and Methods) (Richter et al., 2007). Since live DM cells were normally cultured with DMEM supplemented with 20% FBS, hypertonic conditions were prepared by adding 0.2 M sucrose, and hypotonic conditions were prepared by adding PBS. RI and mitotic DM chromosomal volume were measured by ODT (Fig. 2, Fig. S2). Fig. 2A–C shows representative RI images of live DM cells in metaphase at three different osmotic conditions. It should be noted that the same pseudo-color of the cytosol and chromosome was used for structural clarity and comparison between the conditions. Mitotic chromosomes appeared as large and continuous rod-like RI regions at the center of the mitotic cell. However, many small and dot-like unknown structures were also detected throughout the cytosol. Since such small cytosolic bodies were not detected in the confocal observation of the mitotic chromosome in H2B-mRFP expressed DM cells, these are not likely components of the chromosome, but may be other cytosolic organelles or subcellular bodies. Therefore, small bodies with a similar RI to the chromosome region were excluded from evaluation of chromosome volume prior to comparison (Fig. 1A). Although the mean value of the chromosome volume varied with osmotic conditions, these changes were not significant (Fig. S2). In contrast, confocal images of single DM cells indicated that the mean mitotic chromosome volume was significantly decreased and increased by hypertonic and hypotonic culture, respectively (Fig. 2D, Fig. S3). Mean RI values of mitotic cytosol and chromosomes at each medium condition were evaluated and shown in Fig. 2E. The mean values of RI for the mitotic cytosol and chromosomes in normal DMEM culture conditions were significantly increased by strong hypertonic treatment, while under weak hypotonic conditions, RI values did not decrease significantly compared with normal culture (p > 0.05). The mean RI for the cell membrane did not vary with osmotic stress and was used as a reference. Previous studies showed that osmotic change or molecular crowding induces reorganization of the nuclear structure, as well as noticeable compaction of chromatin in the nucleus (Albiez et al., 2006; Delpire et al., 1985; Finan et al., 2011; Richter et al., 2007; Robbins et al., 1970). To confirm chromatin compaction, the same individual DM cells were analyzed before and after hypertonic or hypotonic treatment (Fig. S4). Fluorescent chromatin in the nuclei of individual DM cells was observed by confocal microscopy before and after treatment. Chromatin was significantly compacted following hypertonic treatment (Fig. S4a, c, e, g, i), but the distribution of chromatin hardly changed with hypotonic treatment (Fig. S4b, d, f, h, j). These findings are in agreement with previous confocal and electron microscopy observations (Richter et al., 2007; Richter et al., 2008). Since the molecular component of chromatin is similar to that of the chromosome, this suggests that compaction of mitotic chromosomes may also be induced by hyperosmotic conditions, and may be accompanied by an increase of molecular density if there is no loss of molecular components in the chromosome under hypertonic conditions (Hancock, 2012).

**Figure 2.**
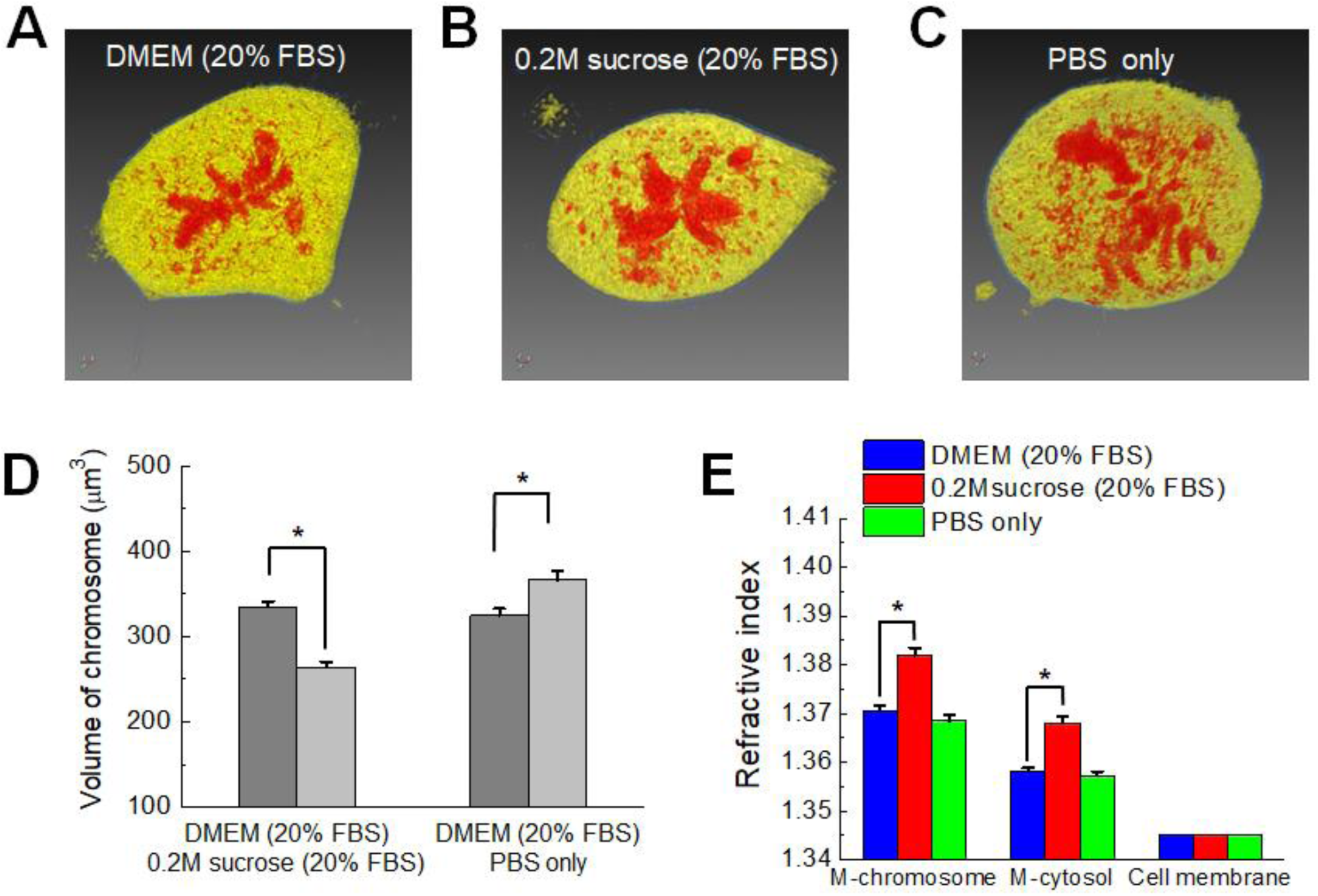
Detection of RI change in chromosomes at different osmotic conditions. (**A**–**C**) Representative RI images of cytosol (yellow), chromosome (red), and plasma membrane (blue) in metaphase of live DM cells under normal (DMEM), hypertonic (0.2 M sucrose), and hypotonic (PBS) conditions. The image of the chromosome does not reflect the true RI value, but only represents the structure of the chromosome. (**D**) Mean chromosome volume under three different conditions (mean ± SEM; n = 25 cells) obtained by 3D confocal imaging of individual DM cells expressing H2B-mRFP. * *p* < 0.05. (**E**) Mean values of RI for chromosomes and cytosol during mitosis at the three different conditions (mean ± SEM; n = 20 cells). The RI value of the cell membrane was not changed and is shown as a reference. M denotes mitotic cell. * *p* < 0.05.

### 3.3 Quantification of the diffusional mobility of mGFP in mitotic chromosomes at different osmotic conditions

Tracking the mobility (i.e., the diffusion coefficient, *D*) of probe molecules is a well-known method used to calculate the local viscosity of fluids and the state of molecular crowding in cells and aqueous solutions (Dix and Verkman, 2008; Pack et al., 2006). As previously demonstrated (Hihara et al., 2012), diffusional mobility of mGFP was quantified in the cytosol and chromosome during mitosis in live DM cells under varying osmotic conditions (Fig. 3). Since the volume of the mitotic chromosome was larger in DM cells than that of other cell lines (Fig. 3A), it was feasible to target and measure the FCS on the mitotic chromosome and cytosol of single DM cells. In agreement with previous findings (Hihara et al., 2012), the fluorescence autocorrelation curves obtained from DM cells fitted well to the single component model of free diffusion, revealing that the diffusional mobility of mGFP in mitotic chromosome was much lower than that in mitotic cytosol (Fig. 3B). The diffusion mobility of mGFP in the cytosol and chromosome changed with osmotic pressure (Fig. 3C, D). Table 2 summarizes the mean *D* values for mGFP in each medium. The average *D* values for mGFP in mitotic cytosol and chromosomes decreased remarkably under hypertonic culture conditions. Although the relative change (<10%) under the hypertonic conditions was smaller than that induced by hypertonic conditions (>40%), the mGFP *D* values were significantly increased under hypotonic cultures. These results demonstrate that mobility of the probe molecule is much more sensitive to changes in the cellular microenvironment induced by osmotic stress. Nevertheless, the FCS results are consistent with the results that showed the RI of chromosomes is influenced by osmotic stress Fig. 2E).

**Table 2.**
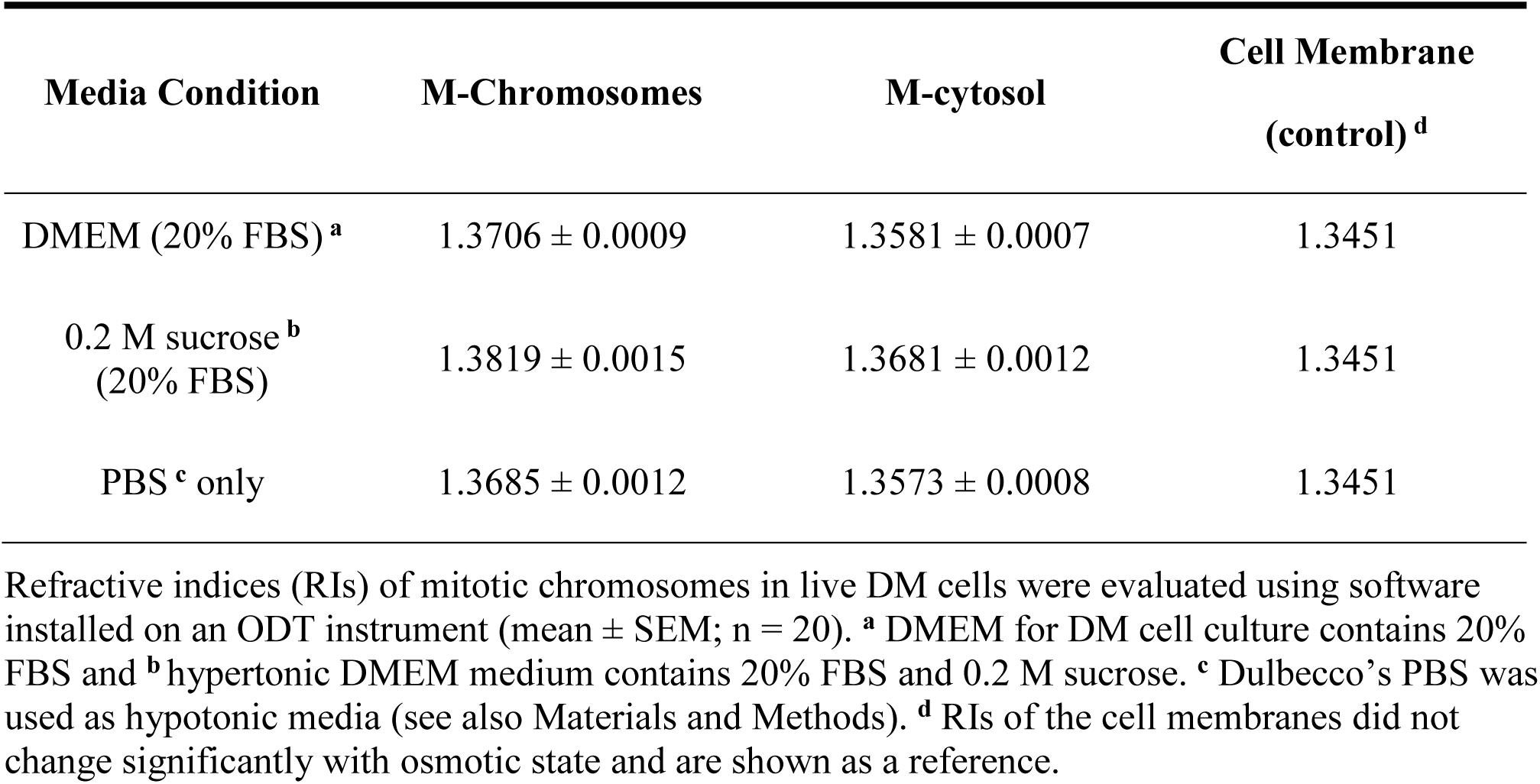
Summary of refractive indices of mitotic chromosomes and cytosol in live DM cells.

| Media Condition | M-Chromosomes | M-cytosol | Cell Membrane<br>(control)<sup>d</sup> |
| --- | --- | --- | --- |
| DMEM (20% FBS)<sup>a</sup> | 1.3706 ± 0.0009 | 1.3581 ± 0.0007 | 1.3451 |
| 0.2 M sucrose<sup>b</sup><br>(20% FBS) | 1.3819 ± 0.0015 | 1.3681 ± 0.0012 | 1.3451 |
| PBS<sup>c</sup> only | 1.3685 ± 0.0012 | 1.3573 ± 0.0008 | 1.3451 |

Refractive indices (RIs) of mitotic chromosomes in live DM cells were evaluated using software installed on an ODT instrument (mean ± SEM; n = 20). <sup>a</sup> DMEM for DM cell culture contains 20% FBS and <sup>b</sup> hypertonic DMEM medium contains 20% FBS and 0.2 M sucrose. <sup>c</sup> Dulbecco's PBS was used as hypotonic media (see also Materials and Methods). <sup>d</sup> RIs of the cell membranes did not change significantly with osmotic state and are shown as a reference.

**Figure 3.**
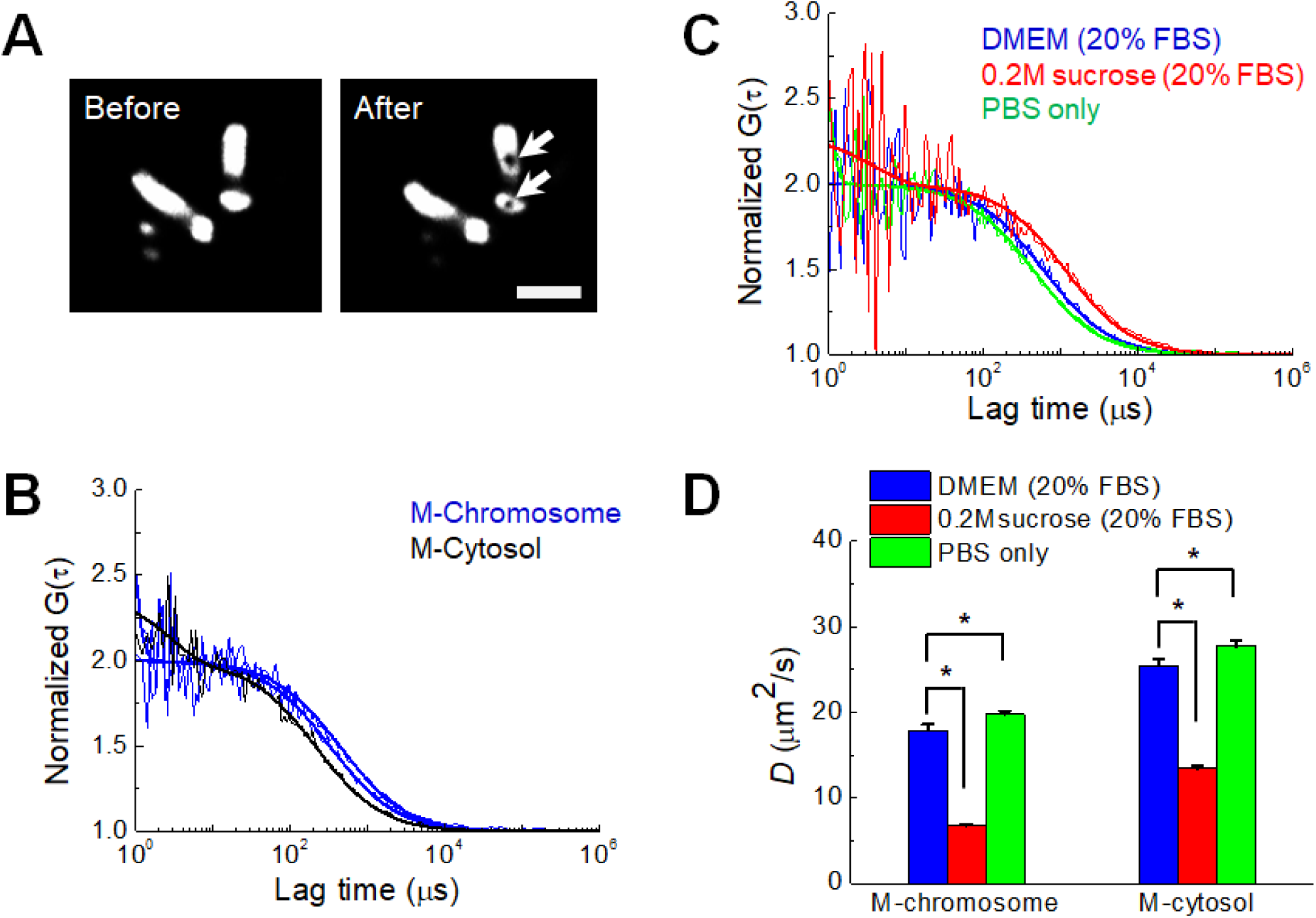
Change in *D* value of mGFP probes in chromosomes at different osmotic conditions. (**A**) Images of mitotic chromosomes (H2B-mRFP) during metaphase of a DM cell before and after two positions of FCS measurement. The positions of FCS measurement in the chromosome are clearly bleached (arrow), verifying the actual measured region. Scale bar represents 5 μm. (**B**) FCS analysis of mGFP expressed in a DM cell in mitosis at normal condition. FAF curves measured in a position of the mitotic cytosol and two positions of mitotic chromosome are shown. (**C**) FAF curves measured in mitotic chromosomes at three different osmotic conditions. For comparison of mobility, all curves in (**B**) and (**C**) were normalized to the same amplitude, *G* (0) = 2. Solid lines indicate fitting of one-component free diffusion model to the results. (**D**) Mean *D* values of mGFP in mitotic chromosome and the cytosol in mitosis at the three different conditions are shown (mean ± SEM; n = 20 cells). * *p* < 0.001.

### 3.4 Relationship between chromosomal RI and diffusion of intrachromosomal mGFP

To quantify the relationship between the RI of the aqueous medium and the diffusion coefficient of fluorescent probe molecules, RIs of water and glycerol–water solutions of different densities were measured. Linear relations between the RI and the molecular density of glycerol solution were obtained at 25°C and 37°C (Fig. 4A, Fig. S5). Previous studies reported that the cellular viscosity of cultured cells is much higher than that of pure water (Dix and Verkman, 2008; Hihara et al., 2012; Pack et al., 2006). The viscosity of solutions with glycerol concentrations from 0% to 75% (w/w) is similar to the fluidic viscosity of the cellular microenvironment. Since the viscosity of the glycerol– water solution is well-known, determination of *D* from FCS measurements of rhodamine 6G (0.479 kD) in glycerol–water solutions provides an experimental relationship between RI and *D* (Fig. 4A–C). From the experimental result and the Stokes–Einstein relationship, theoretical *D* values of mGFP with a molecular weight of 28 kDa in glycerol–water solutions were calculated as shown in Fig. 4D (black) (Pack et al., 2006). The measured mean *D* values (*see also* Table 3) of mGFP in the mitotic cytosol (blue) and DM chromosome (red) shown in Fig. 3 were plotted for comparison with the theoretically calculated *D* values (Fig. 4D). For DMEM and PBS media, the measured *D* values of mGFP in mitotic DM cells were consistent with the theoretical *D* values. Interestingly, the *D* value in DM cell cultured by hypertonic media showed much smaller than the theoretical value. Since hypertonic stress enforces molecular crowding in the cellular components due to the osmotic extraction of water from the cells (Richter et al., 2007), these results suggest that molecular diffusion in the cytosol and chromosome at high osmolality is largely affected by molecular crowding induced by the hypertonic conditions (Golkaram et al., 2016; Park et al., 2015).

**Table 3.**
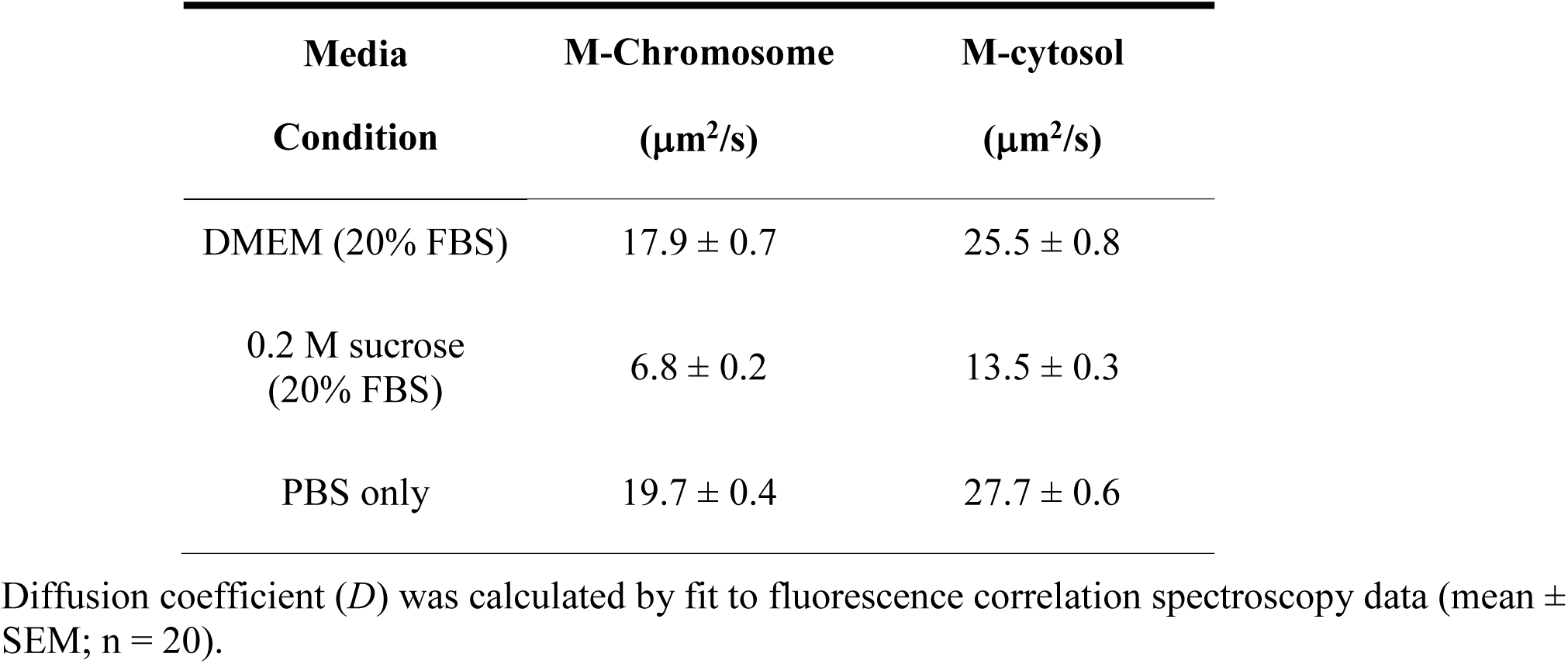
Summary of diffusion coefficient (*D*) of mGFP in mitotic chromosomes and cytosol of live DM cells.

**Figure 4.**
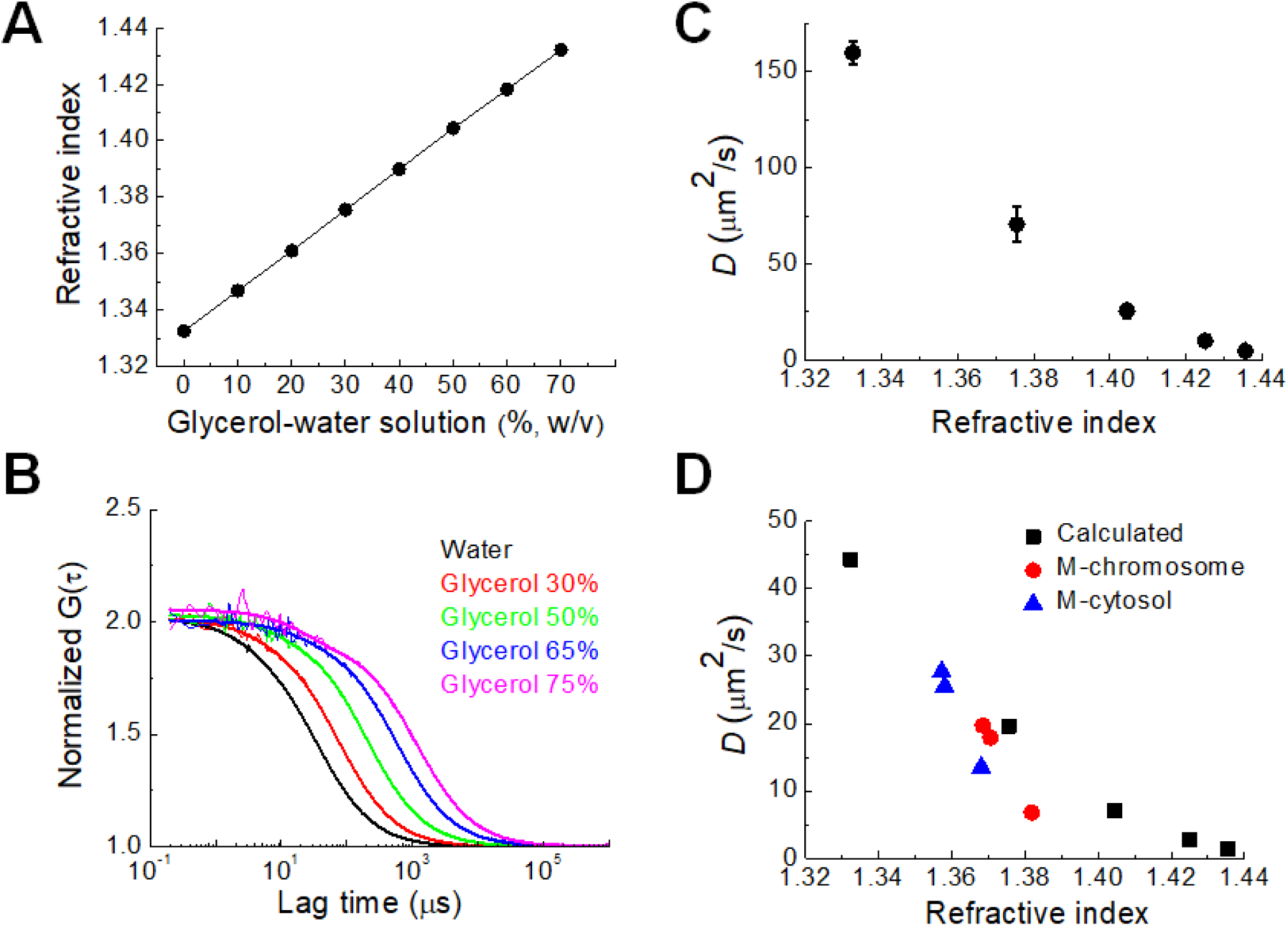
Experimental determination of the relationship between *D* value of rhodamine 6G and RI of medium. (**A**) Measured RI of glycerol–water solution with known glycerol concentration at 25°C (**B**) FAF curves of rhodamine 6G in the glycerol–water solutions. For comparison of mobility, all functions were normalized to the same amplitude, *G* (0) = 2. Bold solid lines indicate fitting of one-component diffusion model to the results. (**C**) Relationship between *D* values of rhodamine 6G in glycerol–water solutions and the RI values in the solutions. (**D**) Plot of calculated *D* value of mGFP in glycerol–water solution with known viscosity versus measured RI (black). Measured *D* values of mGFP in mitotic cytosol (blue) and chromosome (red) at hypotonic, DMEM, and hypertonic culture conditions are shown.

## 4 Conclusion

The present study demonstrates a new method to examine the physicochemical properties of the mitotic chromosome of live cells by combining label-free ODT imaging with complementary confocal microscopy and FCS analysis. The experimental approach of the study is based on the assumption that the RI obtained from ODT analysis is quantitatively related to the diffusion coefficient of the probe molecule from FCS analysis. We verified 3D RI images of mitotic chromosomes in DM cells expressing H2B-mRFP and mGFP by a comparison with 3D confocal images. Finally, we examined the effect of osmotic stress on the RIs of mitotic cells and diffusion coefficient of the mGFP probe molecule. This study successfully demonstrated an inverse relationship between RI and diffusion coefficient in live cells during mitosis. Interestingly, it was found that the chromosome was still accessible to probe molecules that freely diffuse, even though hypertonic stress significantly increased the molecular density of the chromosome. Our results also confirm and extend those of previous studies concerning the influence of osmotic stress on nuclear structure and molecular crowding effect. Our approach provides further insight into the physicochemical properties of chromosomes and other subcellular bodies with a high macromolecular content in live cells.

## 6 Author Contributions

TKK, BWL, FF, and CGP performed the experiments. YGP and KYL performed RI image analysis. TKK, BWL, and CGP collected data and performed data analysis and interpretation. JKK and CGP wrote the manuscript. JKK and CGP supervised and financially supported the study. All authors commented on the manuscript.

## 8 Acknowledgments

We would like to thank Anton Parr Korea Ltd. for the use of their refractometer and their technical support. We also thank Nikon Korea Ltd. and Leica Korea Ltd. for the use of their confocal microscope.

